# Effects of arsenic on the topology and solubility of promyelocytic leukemia (PML)-nuclear bodies

**DOI:** 10.1101/2021.01.13.426540

**Authors:** Seishiro Hirano, Osamu Udagawa

**Affiliations:** Center for Health and Environmental Risk Research, National Institute for Environmental Studies, Japan

**Author notes:** Corresponding author: Seishiro Hirano.

**Keywords:** promyelocytic leukemia-nuclear body (PML-NB), arsenic, small ubiquitin-like modifier (SUMO), ubiquitin, green fluorescence protein (GFP), solubility

## Abstract

Promyelocytic leukemia (PML) proteins are involved in the pathogenesis of acute promyelocytic leukemia (APL). Trivalent arsenic (As^3+^) is known to cure APL by binding to cysteine residues of PML and enhance the degradation of PML-retinoic acid receptor α (RARα), a t(15;17) gene translocation product in APL cells, and restore PML-nuclear bodies (NBs). The size, number, and shape of PML-NBs vary among cell types and during cell division. However, topological changes of PML-NBs in As^3+^-exposed cells have not been well-documented. We report that As^3+^-induced solubility shift underlies rapid SUMOylation of PML and late aggregation of PML-NBs. Most PML-NBs were toroidal and irregular-shaped in *GFPPML*-transduced CHO-K1 and HEK293 cells, respectively. The annular PML-NBs appeared unstable and dissipated into small PML-NBs in HEK cells. Exposure to As^3+^ and antimony (Sb^3+^) greatly reduced the solubility of PML and enhanced SUMOylation within 2 h, and prolonged exposure resulted in PML-NB agglomeration. Exposure to bismuth (Bi^3+^), another Group 15 element, did not induce any of these changes. ML792, a SUMO activation inhibitor, reduced the number of PML-NBs and increased the size of the NBs, but had little effect on the As^3+^-induced solubility change of PML. The results show that SUMOylation regulates the dynamics of PML-NBs but does not contribute to the As^3+^-induced solubility change of PML.

## Introduction

Promyelocytic leukemia proteins (PMLs) are proapoptotic molecules involved in cell proliferation, senescence, and tumor suppression (1). The human PML family comprises six known nuclear isotypes (I-VI) and one cytosolic isotype (VII) (2–4). The life-time of PML ranges from several minutes to one hour, whereas other PML nuclear body (PML-NB) client proteins such as Sp100 and Daxx range from several seconds to one minute (5), which led to the notion that PML is a scaffold protein for PML-NBs. Sp100 is not essential for the maintenance of PML-NBs in human embryonic NT2 cells and the C-terminal region of Daxx is sufficient for interaction with PML (6). The mutation of three small ubiquitin-like modifier (SUMO) conjugation sites of PML (K65, K160, and K490) did not affect the formation of PML-NBs, although this mutant could recruit neither Sp100 nor Daxx (7). The SUMOylation of PML is thought to stabilize and promote the assembly of PML-NBs, since free PML tends to disperse throughout the nucleoplasm and SUMO1-conjugated PML localizes to PML-NBs (8). PML-NBs are structurally and positionally stable in interphase and appear to associate with euchromatin via SUMO molecules (9).

PML is a member of RBCC/tripartite motif (TRIM) family. Cysteines in both B_1_ and B_2_ domains are required for PML-NB formation (10). As^3+^ binds to cysteine residues of the RING and B_2_ domains (11) and the core region of RING is requisite for the solubility change and SUMOylation of PML in response to exposure to As^3+^ (12). C_212_ in the B_2_ domain is required for As^3+^-induced degradation of PML-RARα, a t(15;17) gene translocation product (13). The subdomain comprising F_52_Q_53_F_54_, and L_73_ in the RING motif (aa 55-99) are unique to PML and are absent in other TRIM family members. The unique subdomain appears to play a role in PML tetramer and the following subsequent PML-NB formation, and also in As^3+^-induced SUMOylation of PML (14). Together, intact zinc finger domains of RBCC and the unique RING motif of PML are all necessary for PML-NB formation and biochemical responses of PML to As^3+^.

PML-RARα functions as a dominant negative form and causes acute promyelocytic leukemia (APL) (15,16). Both As^3+^ and all-trans retinoic acid (ATRA) have been used to treat APL (17,18). As^3+^-binding to PML renders PML-RARα susceptible to ubiquitin-mediated degradation by RNF4 which non-covalently binds to SUMOylated PML via SUMO-interacting motifs (SIMs) (3,19).

Canonical PML-NBs are present in the nucleus as dense granular bodies or hollow toroid-like oblate (doughnut shaped) spheres (20), and the shape and size of PML-NBs vary depending on the cell type (21). Non-spherical forms of PML-NBs have also been reported. Fiber-like PML-NBs are formed along with doughnut-like PML-NBs in the nuclei of IDH4 cells, human fibroblasts immortalized with the dexamethasone-inducible SV40 (22). Nuclear envelope-associated linear and rod-like PML structures reportedly arise transiently during the very early transitions of human ES cells towards cell-type commitment (23).

Although PML-NBs are stable in interphase nuclei, they aggregate to form mitotic accumulation of PML proteins (MAPPs) after nuclear membrane breakdown. MAPPs associate with nuclear pore components in daughter cells after mitosis. At this stage the aggregated PML bodies are called cytoplasmic assemblies of PML and nucleoporins (CyPNs) and reside in peri-nuclear regions (24,25). MAPPs are inherited asymmetrically in daughter HaCaT cells after mitosis (26), and daughter cells that receive the majority of PML-NBs exhibit increased stemness in primary human keratinocytes (25). Since PML-NBs feature phase-separated protein condensates in the nucleus (27,28), it is of interest to investigate how their biochemical/biophysical properties change upon cell division and exposure to As^3+^.

In the present work, we performed a topological characterization of PML-NBs using CHO-K1 and HEK293 cells stably transduced with GFP-conjugated PML-VI. Then, we investigated how As^3+^ and other trivalent Group15 elements affect the biochemical characteristics of PML and the dynamics of PML-NBs. We also studied effects of As^3+^ on PML and PML-NBs in the presence of a SUMOylation inhibitor.

## Results

### Topology and dynamics of PML-NBs

Most interphase PML-NBs appeared to group into large and small toroid-like forms in CHOGFPPML cells (Fig. 1A). The small PML-NBs often accompany peri-nuclear large toroidal PML structures (the box with white line in Fig, 1A), and the volume ratio of the large to small toroids was calculated to be 8.4, assuming that those toroids were symmetrical doughnuts. We next studied time-course changes in the shapes of PML-NBs by confocal microscopy using the z-stack mode. Figure 1B shows that PML aggregates (MAPPs) appeared after nuclear membrane breakdown in dividing CHOGFPPML cells (0:00:00). Another large toroidal PML aggregate appeared quickly in the peri-nuclear region via the condensation of vague PML speckles (arrow) in 87 sec (from 0:11:55 to 0:13:22) as cytokinesis proceeded (See also Supporting Fig, 1A for freshly formed peri-nuclear PML structures). Uneven partitioning of peri-nuclear PML aggregates was observed in dividing untreated (Supporting Fig. 1A) and As^3+^-exposed CHOGFPPML cells (Supporting Fig. 1B). Small nascent PML-NBs were formed as the daughter cells spread. Contrary to CHOGFPPML cells, annular PML-NBs were very rare and most PML-NBs in HEKGFPPML cells appeared as irregular agglomerates. PML-NBs in HEKGFPPML cells were grouped into large and small dots (Fig. 2A). The sizes of PML-NBs in HEKGFPPML cells were estimated as the mean of the longest and shortest axes. We found a HEKGFPPML cell harboring several toroidal PML-NBs (Fig. 2B). The sizes of these toroidal PML-NBs were comparable to those of CHOGFPPML cells. These toroidal PML-NBs in HEKGFPPML cells were, however, gradually dissipated and transformed into small PML-NBs in 1.5 h (boxed areas), suggesting that the toroidal PML-NBs in HEKGFPPML cells are less sable than those in CHOGFPPML cells.

**Fig. 1.**
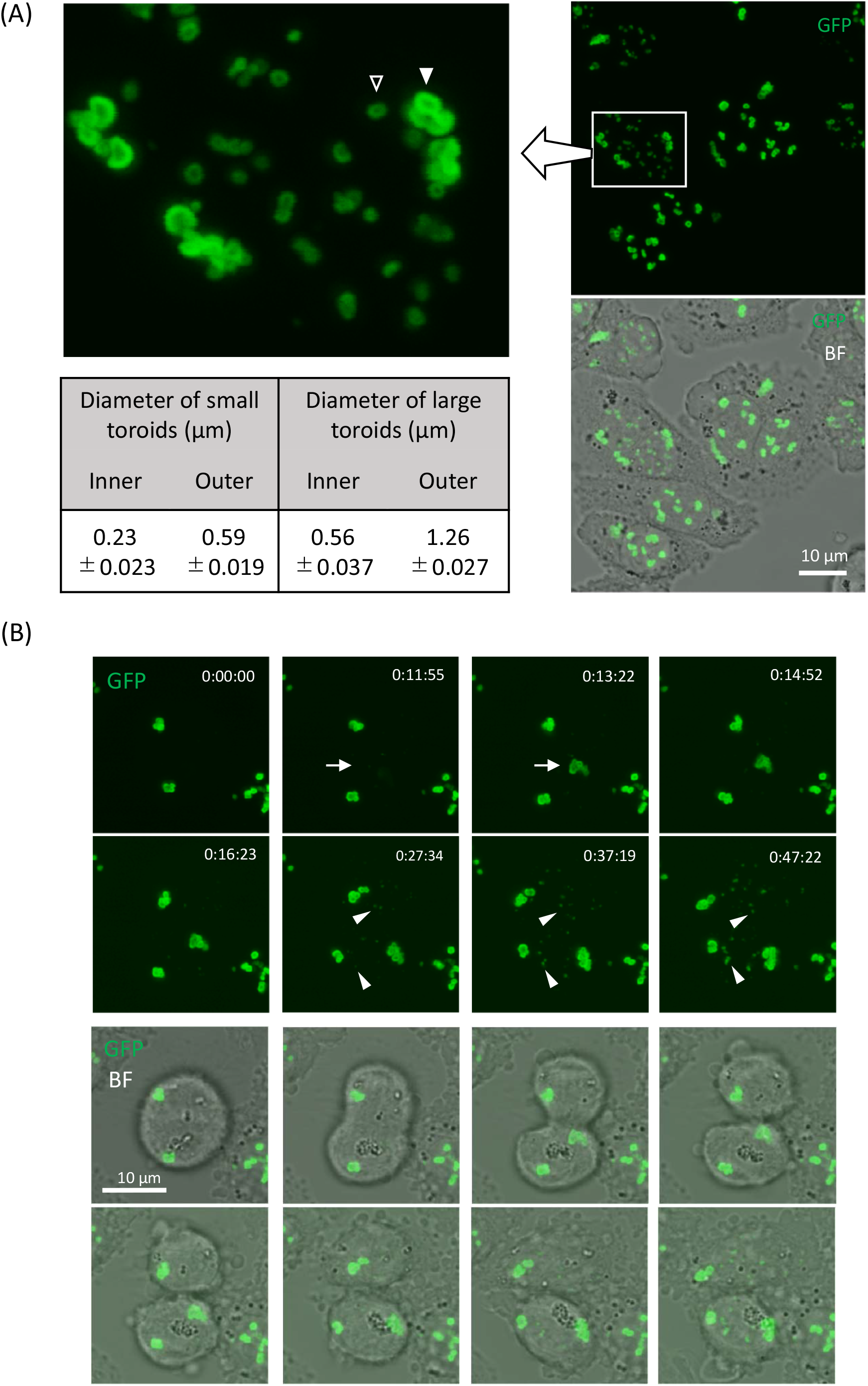
The structure and size of PML-NBs and peri-nuclear PML structures (A) and formation of peri-nuclear PML structures and small nascent PML-NBs in CHOGFPPML (B). GFP live images were obtained by confocal laser microscopy in z-stacking mode. (A) PML-NBs occurred as either small or large toroids in CHOGFPPML cells. The white boxed area was enlarged to visualize the shapes of PML-NBs and peri-nuclear PML structures clearly. The volume of each toroid was calculated from the inner and outer diameters, assuming that the toroid has a doughnut shape. The volume ratio of the large toroid (closed triangle) to the small toroid (open triangle) was calculated to be 8.4. Data are presented as means ± SEM (N=7). (B) Time-lapsed images were captured during cell division and spreading. The upper 8 panels show GFP images and the lower panels are overlaid with corresponding bright field (BF) images. The time counter is shown in the right upper corner of each GFP panel. Arrows show that *de novo* appearance of the large peri-nuclear PML structure; The accretion of GFP speckles began at 0:11:55 and the large toroidal PML structure formed clearly after 1 min and 27 sec (at 0:13:22). Arrowheads show small nascent PML-NBs that appeared in daughter cells after cell division.

**Fig. 2.**
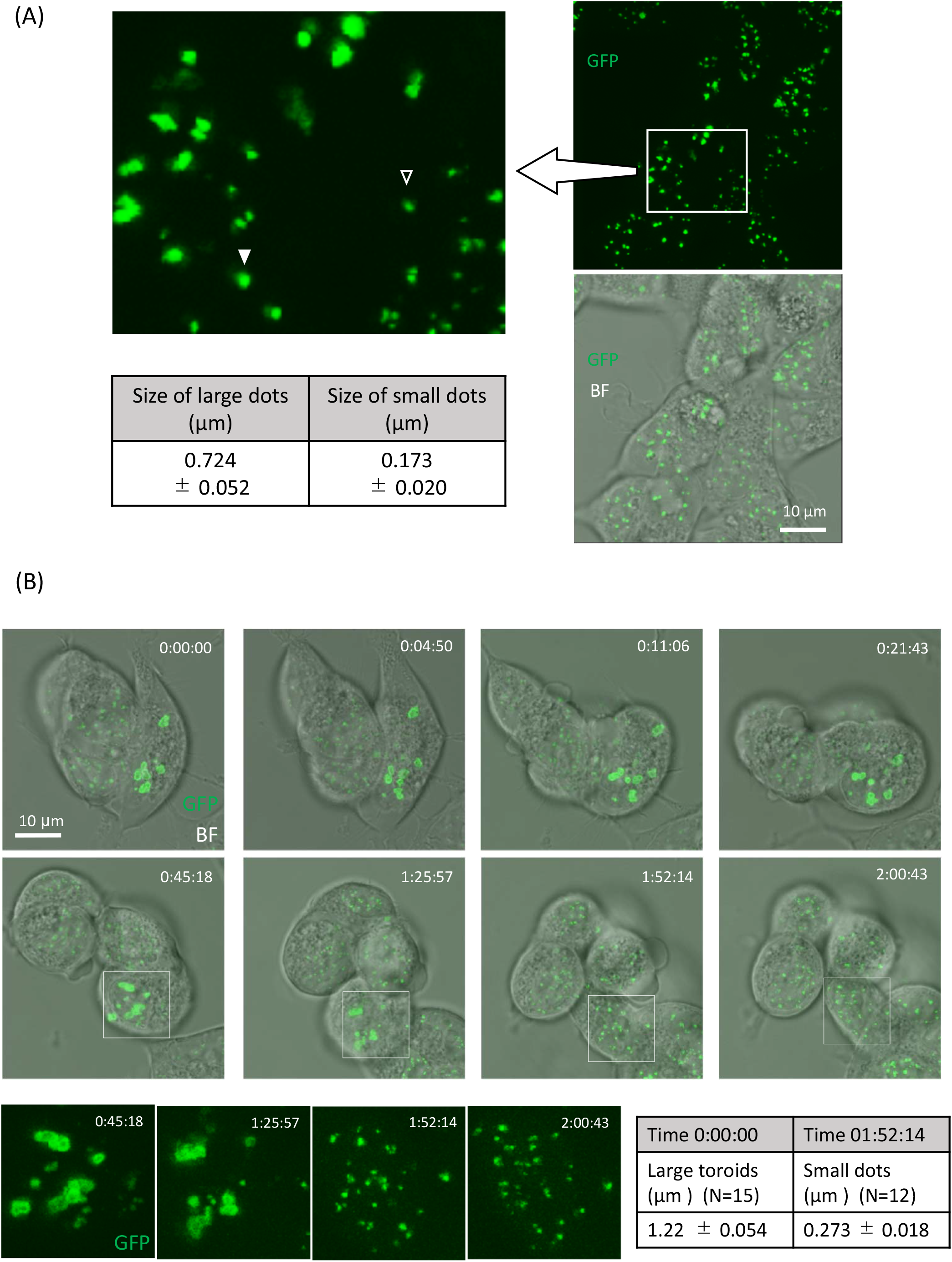
The structure and size of PML-NBs (A) and conversion of large toroidal PML-NBs into small PML-NBs in HEKGFPPML cells (B). GFP live images were obtained by confocal laser microscopy in z-stacking mode. (A) PML-NBs occurred either as small dots (open triangle) or irregular large dots (closed triangle) in HEKGFPPML cells. The sizes of these dots were evaluated by the average of the major and minor axes, assuming that each dot was oval. Data are presented as means ± SEM (N=10). (B) Time-lapsed images of HEKGFPPML cells were captured. The time counter is shown in the right upper corner of each GFP-BF overlaid panel. GFP fluorescence images in the boxed areas were enlarged and are shown in the bottom 4 panels. The toroid-like PML-NBs dissipated or dissolved (1:25:57), then were transformed into smaller PML-NBs (1:52:14) in non-dividing cells. Sizes of the large toroids (0:00:00) and the small dots (01:52:14) were evaluated by the average of the major and minor axes, assuming that each toroid or dot was oval.

### Effects of As^3+^, Sb^3+^, and Bi^3+^ on PML and intranuclear distribution of PML-NBs

This experiment was performed to study the specificity of As^3+^ as a modifier of intranuclear dynamics of PML-NBs. Both As^3+^ and Sb^3+^ are metalloids, while Bi^3+^ is a heavy metal belonging to Group 15 of the elements. Bi^3+^ was slightly less cytotoxic than either As^3+^ or Sb^3+^ in HEKGFPPML cells (Fig. 3A). Exposure to 3 μM As^3+^ or Sb^3+^ for 2 h induced an overt protein solubility change and converted cold RIPA-soluble GFPPML into the RIPA-insoluble form, and SUMOylated GFPPML. However, these biochemical changes to GFPPML were not observed in Bi^3+^-exposed cells (Fig. 3B). Contrary to the explicit biochemical changes in GFPPML, the microscopically observed size, number, and intranuclear distribution of PML-NBs did not remarkably change in the 2-h treatment with either As^3+^ or Sb^3+^. However, PML-NBs agglomerated in the nucleus of CHOGFPPML cells 24 h after exposure to 1-3 μM As^3+^ or Sb^3+^. Exposure to Bi^3+^ did not cause these microscopic changes (Fig. 4).

**Fig. 3.**
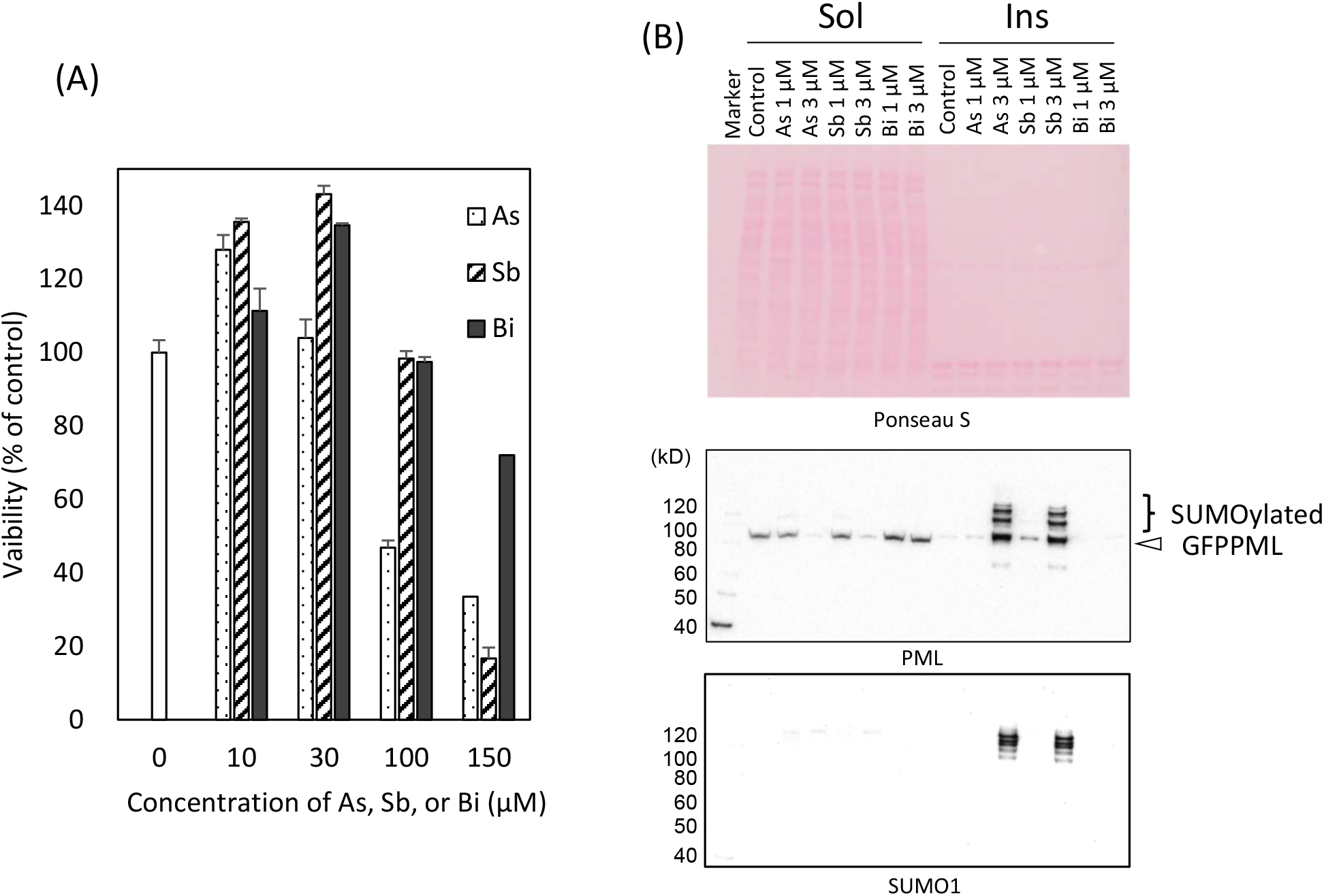
Cytotoxicity (A), and solubility changes and SUMOylation of PML (B) in response to trivalent arsenic (As), antimony (Sb), or bismuth (Bi) ions in CHOGFPPML cells. (A) The cells were exposed to various concentrations of As^3+^, Sb^3+^, or Bi^3+^ for 24 h. As^3+^ was significantly more cytotoxic than Sb^3+^ and Bi^3+^ in CHOGFPPML cells as analyzed by two-way ANOVA followed by Tukey’s multiple comparison. Data are presented as means ± SEM of quadruplicated wells. (B) The solubility of PML in cold RIPA buffer decreased in response to 2 h-exposure to 1–3 μM As^3+^ or Sb^3+^. GFPPML was SUMOylated (SUMO1) following exposure to either 3 μM As^3+^ or Sb^3+^. These biochemical changes were not observed in Bi^3+^-exposed cells. The major cold RIPA-insoluble proteins observed by Ponseau S-staining were histones. ‘Sol’ and ‘Ins’ denote the RIPA-soluble and -insoluble fractions, respectively.

**Fig. 4.**
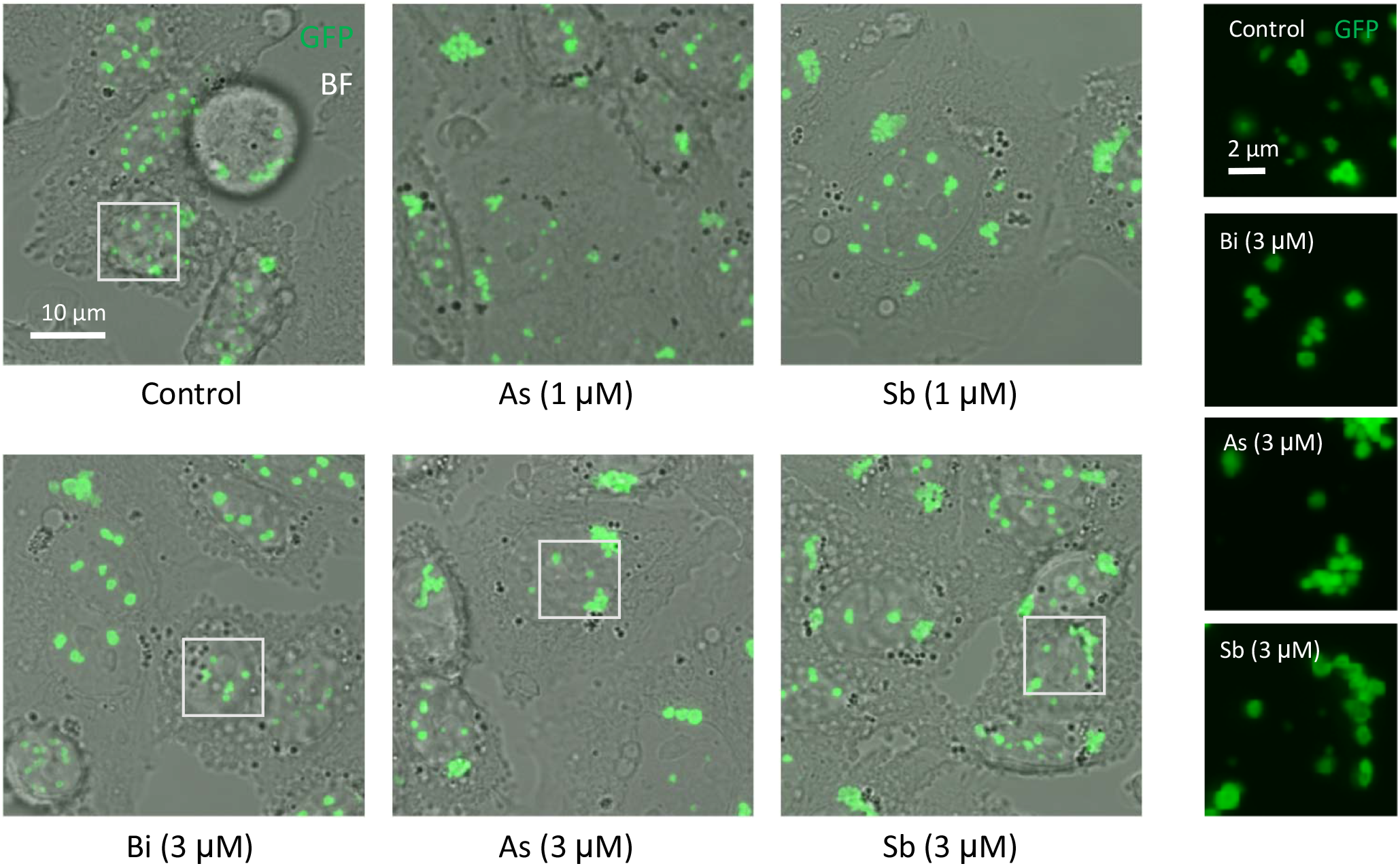
Agglomeration of PML-NBs in response to As^3+^ or Sb^3+^ in CHOGFPPML cells. The cells were exposed to 1 or 3 μM As^3+^, Sb^3+^, or Bi^3+^ for up to 24 h. Contrary to the dramatic solubility change and SUMOylation of PML in response to As^3+^ and Sb^3+^, the intranuclear distribution of PML-NBs was not explicitly changed by 2 h-exposure to As^3+^ or Sb^3+^. After exposure to As^3+^ or Sb^3+^ (1 and 3 μM) for 24 h, however, agglomeration of PML-NBs was observed, while most PML-NBs were evenly distributed in Bi^3+^-exposed and untreated cells. GFP images within the white boxes were enlarged and are shown in the right panels..

We next examined a possibility that As3+ and Sb3+ inhibited the proteasome activity and SUMOylated PML proteins were not eliminated efficiently by ubiquitin-proteasome system (UPS). The 20S proteasome activity was inhibited slightly by 30 μM As^3+^, Sb^3+^, Bi^3+^, and Cd^2+^. However, Bi^3+^ inhibited more efficiently than As^3+^ and Sb^3+^ (Supporting Fig. 2), suggesting that the degradation of SUMOylated PML by UPS was not involved in the different SUMOylation levels of PML proteins between Bi^3+^-exposed and As^3+^- or Sb^3+^-exposed cells. The As^3+^-induced solubility shift and SUMOylation of PML, and aggregation of PML-NBs are not irreversible phenomena, because removal of As^3+^ from the culture medium reverted those As^3+^-induced changes in CHOGFPPML cells (Fig. 5) and in HEKGFPPML cells (Supporting Fig. 3).

**Fig. 5.**
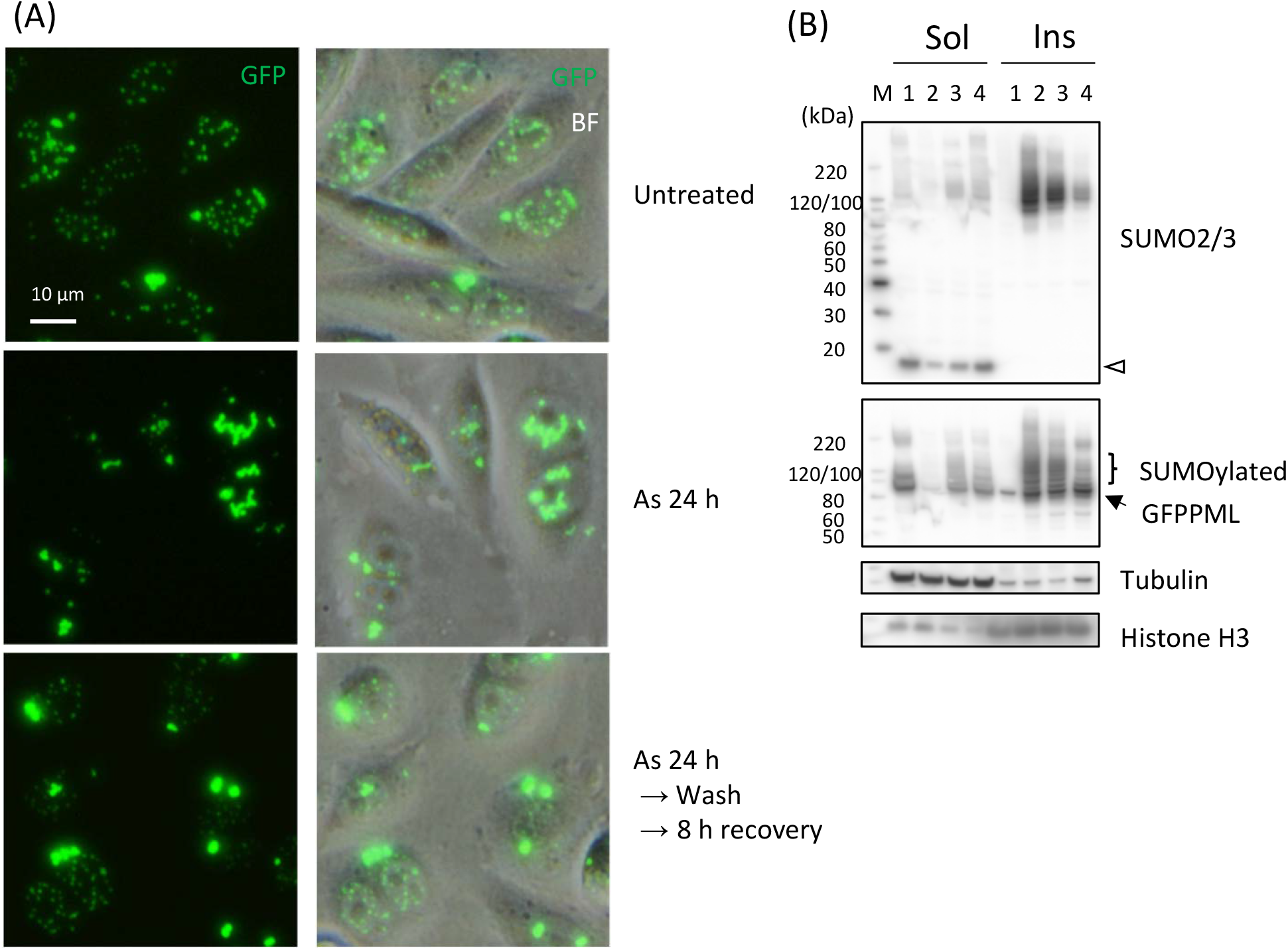
Recovery of nuclear distribution of PML-NBs (A), and GFPPML and SUMO2/3 proteins in the RIPA-soluble fraction (Sol) (B) from a 24 h-exposure to As^3+^. (A) The cells were exposed to 3 μM As^3+^ for 24 h. The cells were washed and further cultured in As^3+^-free culture medium for 8 h. The nuclear distribution of GFPPML was observed by fluorescent microscopy. (B) The As^3+^-exposed cells were lysed with the RIPA buffer immediately or washed and cultured in As^3+^-free culture medium further for 8 or 24 h. 1, untreated; 2, 24 h exposure to As^3+^; 3, 24 h exposure to As^3+^and 8 h recovery in As^3+^-free culture medium; 4, 24 h exposure to As^3+^and 24 h recovery in As^3+^-free culture medium. The unconjugated GFPPML and GFPPML conjugated with SUMO2/3 in the RIPA-insoluble fraction (Ins) decreased during the culture in As^3+^-free culture medium. An open arrowhead indicates SUMO2/3 monomers.

### SUMOylation and Ubiquitination of PML

PML proteins are modified by SUMOylation and subsequent ubiquitination by RNF4 E3 ubiquitin ligase in As^3+^-exposed cells (19,29). We investigated whether post-translational modification of GFPPML with SUMO and ubiquitin molecules occurs while GFPPMLs are still soluble in cold RIPA. Figure 6 shows that exposure to As^3+^ converted GFPPML from the RIPA-soluble to the RIPA-insoluble form, and SUMOylated GFPPML remained in the RIPA-soluble fraction at 1 h post-exposure. Accordingly, immunoprecipitation was performed using the RIPA-soluble fraction with magnetic bead-tagged anti-GFP antibody using the clear supernatants of RIPA lysates obtained from untreated and As^3+^-exposed CHOGFPPML cells (Fig. 7A) and HEKGFPPML cells (Fig. 7B). Modification on GFPPML with SUMO2/3 and SUMO1, and ubiquitination, were clearly observed in these cells after 1 h-exposure to As^3+^. It should be noted that the amount of SUMOylated GFPPML immunoprecipitated in the untreated cells was less than that of As^3+^-exposed cells.

**Fig. 6.**
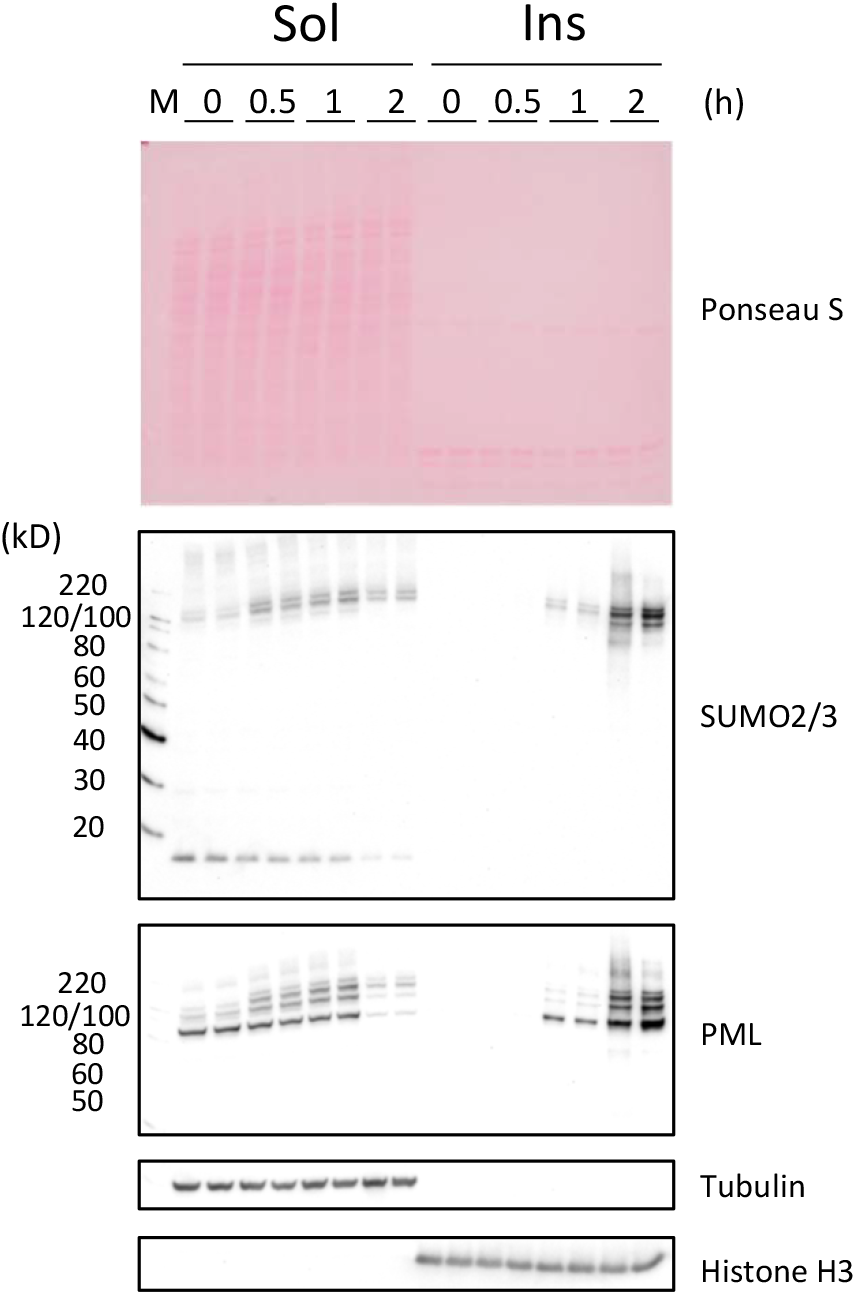
Time-course changes in the solubility and SUMOylation of PML following exposure to 3 μM As^3+^ in CHOGFPPML cells. The cells were exposed to 3 μM As^3+^ for 0.5, 1, and 2 h. SUMOylation of PML was observed in the RIPA-soluble fraction (Sol) after 0.5 h-exposure. The solubility of unmodified PML decreased after 1 h, and most of the PML and SUMOylated PML was recovered in the RIPA-insoluble fraction (Ins) after 2 h-exposure to As^3+^. An open arrowhead indicates SUMO2/3 monomers.

**Fig. 7.**
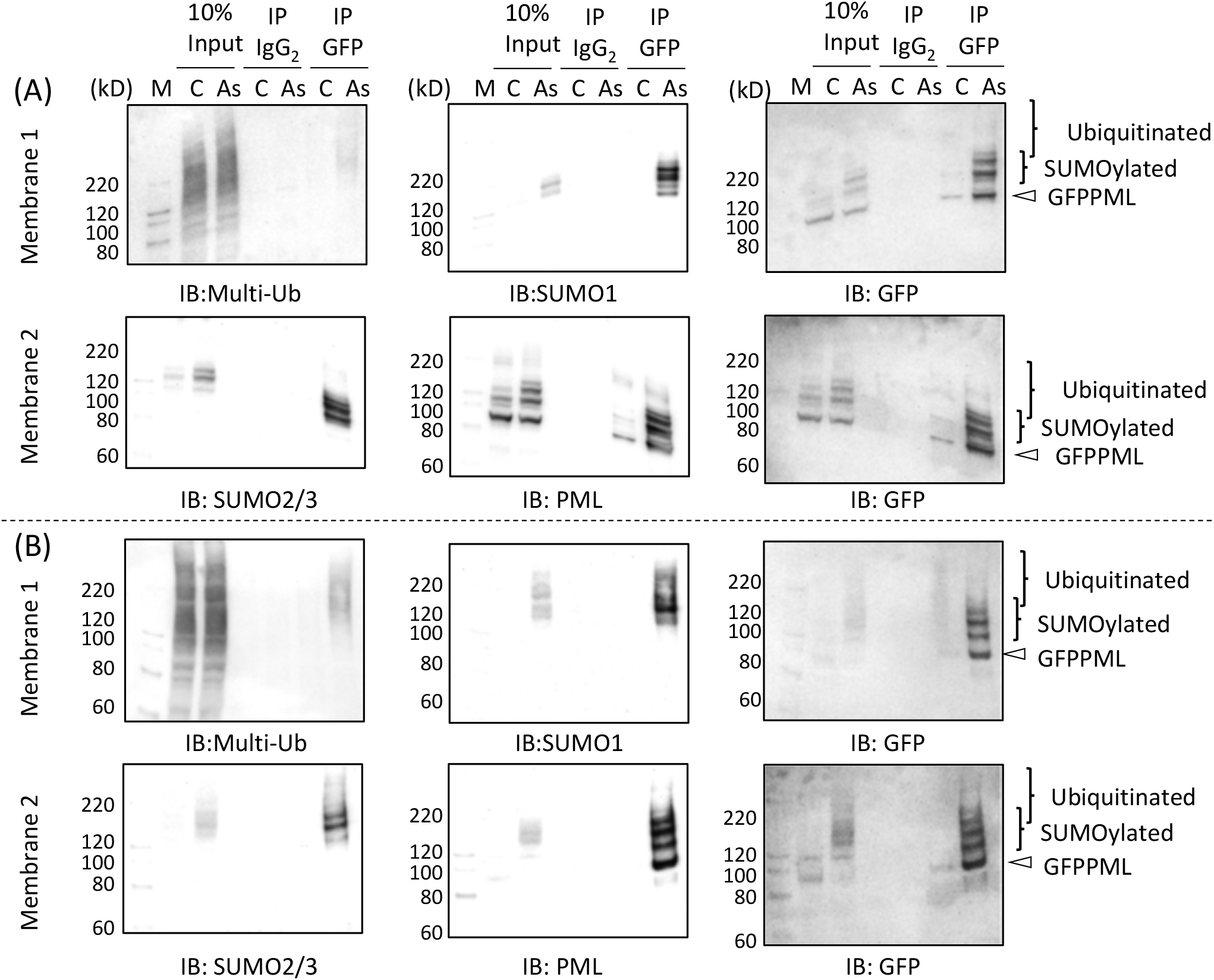
Immunoprecipitation analyses to detect the SUMOylation and ubiquitination of PML in CHOGFPPML (A) and HEKGFPPML cells (B). The cells were exposed to 3 μM As^3+^ (As) or left untreated (C) for 1 h and the RIPA-soluble fractions were collected. The samples were centrifuged again to obtained clear supernatants and subjected to immunoprecipitation with anti-GFP antibody-magnetic beads. The immunoprecipitates were resolved by SDS-PAGE and subsequent western blotting. The membranes were sequentially probed with anti-multi-Ub, anti-SUMO1, and anti-GFP antibodies (membrane 1) or with anti-SUMO2/3, anti-PML, and anti-GFP antibodies (membrane 2).

### Effects of SUMOylation and ubiquitination inhibitors on PML-NBs

Recently, TAK243 and ML792 were shown to effectively inhibit the ubiquitination and SUMOylation of PML, respectively (30). Exposure to 20 μM ML792 for up to 8 h did not significantly change the viability of HEKGFPPML cells, although longer exposure slightly reduced the number of viable cells compared to untreated cells (Supporting Fig. 4A). TAK243 was cytotoxic at a concentration of 10 μM and cells started to shrink after 5 h culture with TAK243. It is noteworthy that ML792 reduced the cytotoxicity of TAK243, suggesting that SUMOylation may enhance the adverse effects caused by the accumulation of ubiquitin-free and improperly folded proteins (Supporting Fig. 4B).

Inhibiting SUMOylation with ML792 reduced the number of PML-NBs and reciprocally increased their sizes in both CHOGFPPML (Fig. 8A) and HEKGFPPML cells (Fig. 8B), although the difference in size was not statistically significant in CHOGFPPML cells. The number of PML-NBs was slightly reduced by TAK243 in CHOGFPPML cells but not in HEKGFPPML cells. These results indicated that the SUMOylation of PML is important to maintain the number and size of PML-NBs. Figure 9 shows that TAK243 completely inhibited the ubiquitination of RIPA-soluble proteins in both CHOGFPPML (Fig. 9A) and HEKGFPPML cells (Fig. 9B). In the RIPA-insoluble fraction, the major anti-multi-ubiquitin reactive protein (24 kD) that disappeared upon TAK243 treatment is probably mono-ubiquitinated H2A and/or H2B (31). SUMOylation of GFPPML with either SUMO1 or SUMO2/3 was inhibited by ML792. SUMO2/3 monomers, present in the RIPA-soluble fraction, essentially disappeared in CHOGFPPML (Fig. 9A) and were reduced in HEKGFPPML cells (Fig. 9B) after 2 h-exposure to As^3+^. However, SUMO2/3 monomers were recovered to the original amount in ML792-pre-treated and As^3+^-exposed cells. Exposure to As^3+^ caused a solubility shift and the SUMOylation of GFPPML. Although ML792 inhibited SUMOylation, non-SUMOylated GFPPML was recovered in the RIPA-insoluble fraction after exposure to As^3+^, suggesting that SUMOylation is not the cause but rather the consequence of the solubility change of GFPPML. Alternatively, the solubility shift and SUMOylation of GFPPML may occur independently in response to As^3+^. UBA2, a SUMO-activating enzyme, was recovered in the RIPA-soluble fraction, regardless of exposure to As^3+^.

**Fig. 8.**
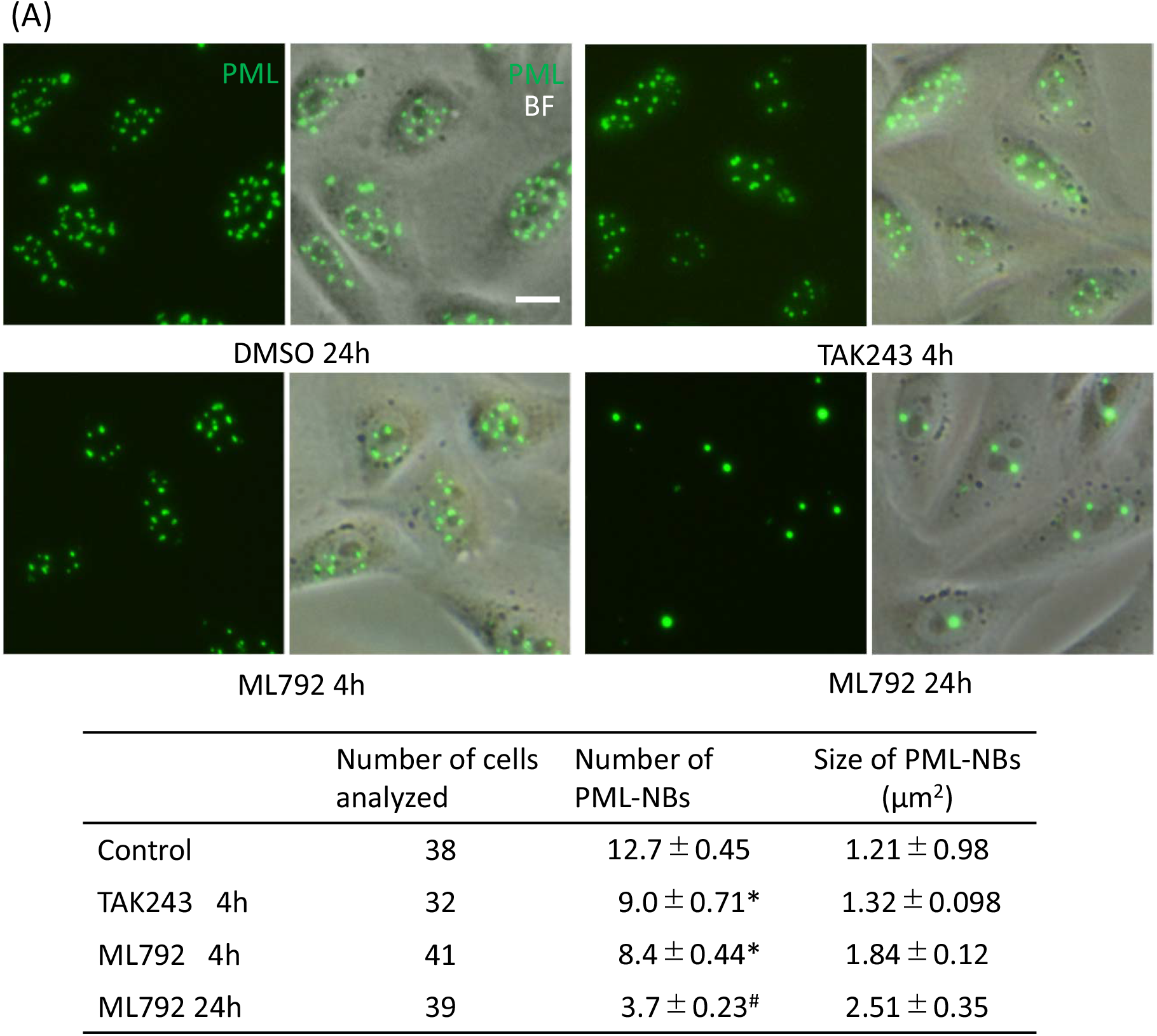
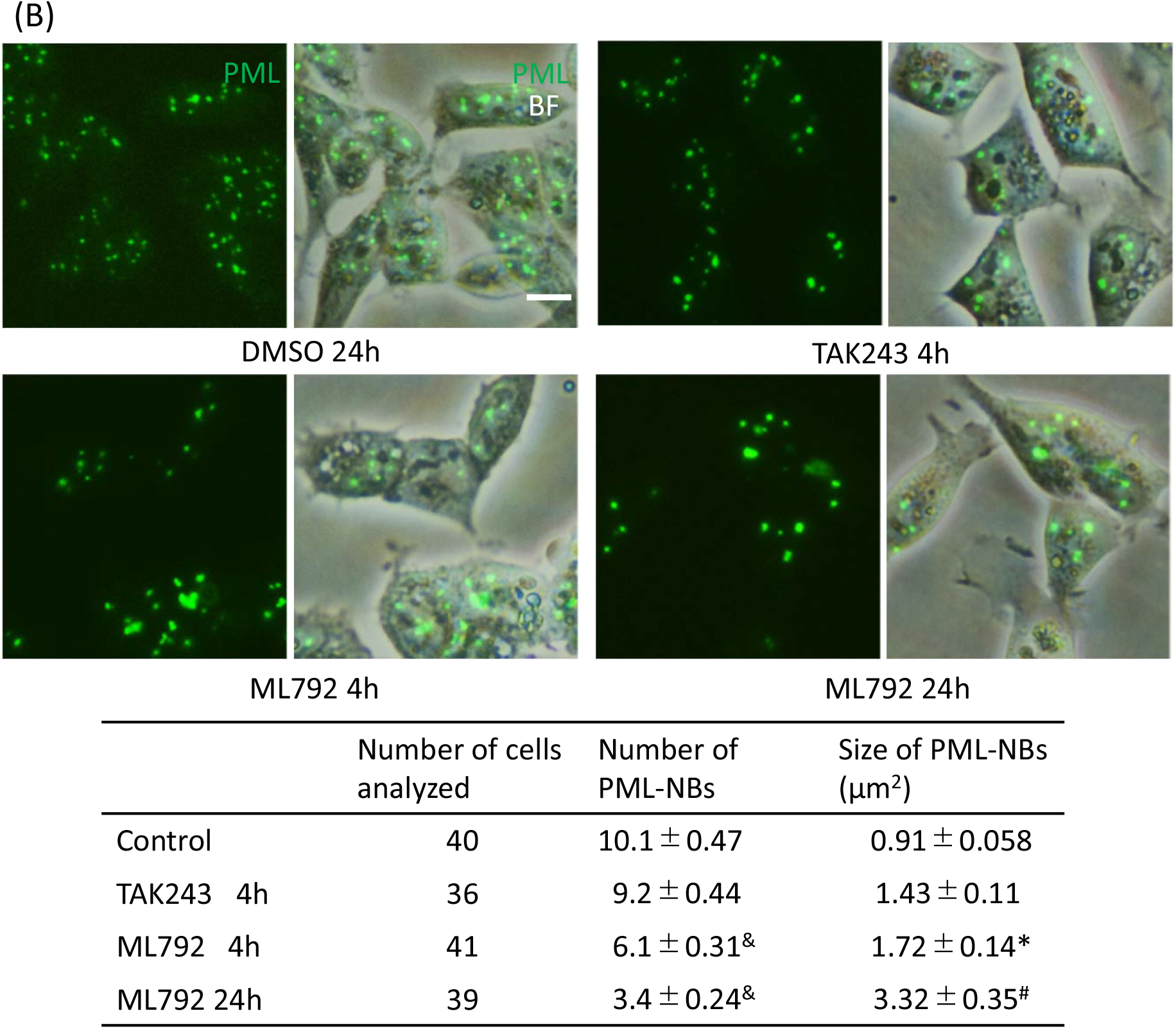
Effects of TAK243 (ubiquitination inhibitor) and ML792 (SUMOylation inhibitor) on the number and size of PML-NBs in CHOGFPPML (A) and HEKGFPPML cells (B). The cells were cultured in the presence of 0.1% DMSO (control), 10 μM TAK243, or 20 μM ML792 for 4 or 24 h. The number of PML-NBs in both cell types decreased and their sizes increased significantly by treatment with ML792. TAK243 was less effective than ML792 in decreasing the number of PML-NBs, and the cells looked shrunken and slightly damaged after 6 h of culture with 10 μM TAK243. Scale bar = 10 μm

**Fig. 9.**
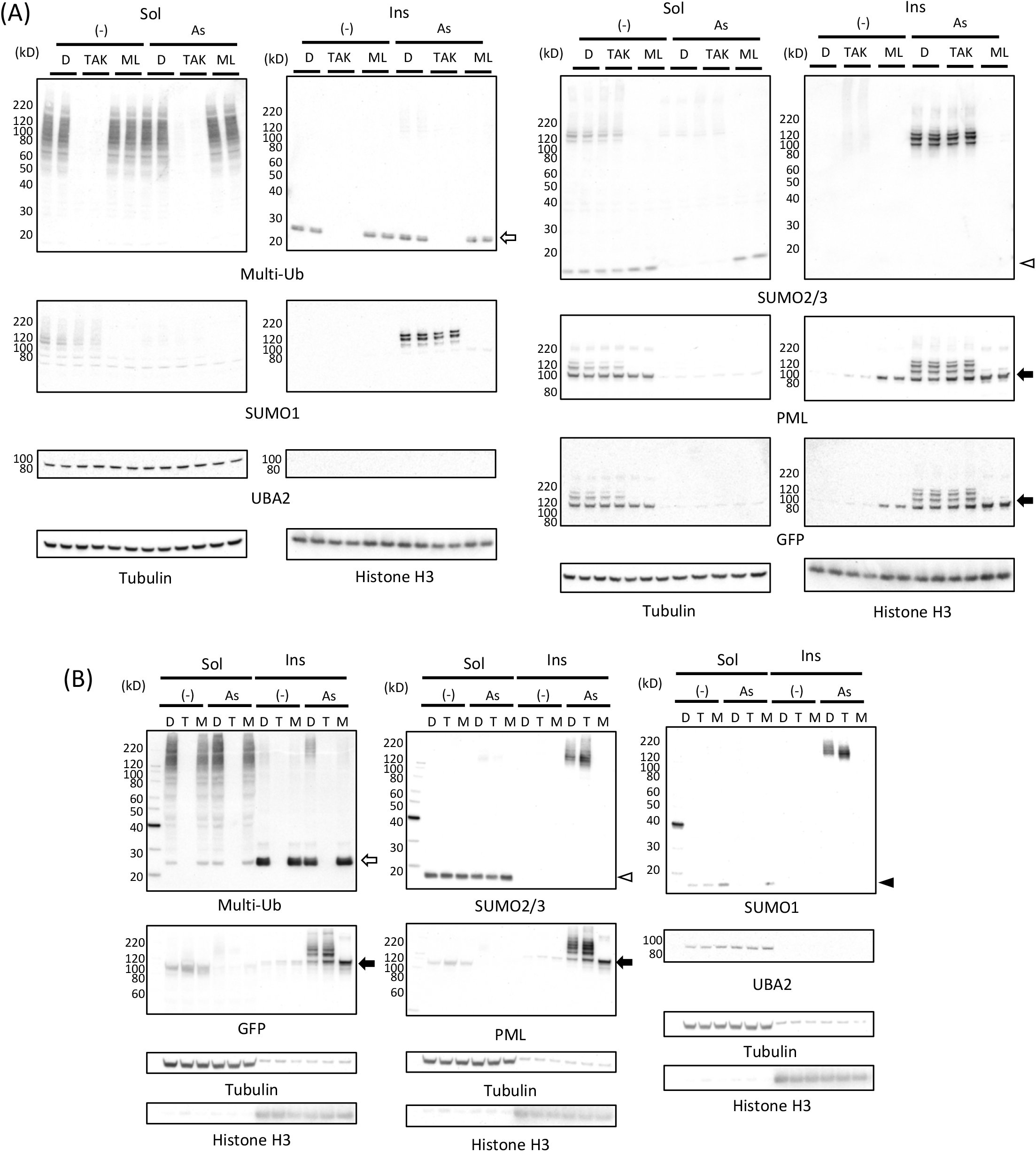
Effects of TAK243 and ML792 on As^3+^-induced biochemical changes of PML in CHOGFPPML (A) and HEKGFPPML cells (B). The cells were pre-treated with 0.1% DMSO, 10 μM TAK243, or 20 μM ML792 for 3 h, then further treated with 3 μM As^3+^ or left untreated for 2 h. The RIPA-soluble and -insoluble fractions were prepared as described previously and the proteins were resolved by SDS-PAGE, followed by western blot. For CHOGFPPML cells, samples following each treatment were obtained from 2 separate wells. The membranes were probed sequentially with the indicated antibodies, then reprobed with a mixture of HRP-tagged anti-tubulin and HRP-tagged anti-histone H3. Open arrow, monoubiquitinated histone; Closed arrow, GFPPML; Open arrowhead, SUMO2/3 monomers; Closed arrowhead, SUMO1 monomers.

Finally, we investigated the intracellular distribution of SUMO2/3 when SUMOylation was inhibited. CHOGFPPML cells were pre-cultured in the presence or absence of ML792 and exposed to As^3+^ or left untreated for 2 h before immunostaining with anti-SUMO2/3. It is of interest to note that the ratio of SUMO2/3 to PML of the peri-nuclear PML structures was lower than that of PML-NBs regardless of As^3+^-exposure in the absence of ML792 (arrows, Fig. 10). However, the SUMO/PML ratio was almost the same between peri-nuclear PML structures and PML-NBs when the cells were pre-treated with ML792. Since some peri-nuclear PML structures are freshly formed by accretion of small PML speckles during cell division (Fig. 1B and Supporting Fig. 1A), it is reasonable to suppose that ML792 disturbs the intranuclear restoration of SUMO molecules.

**Fig. 10.**
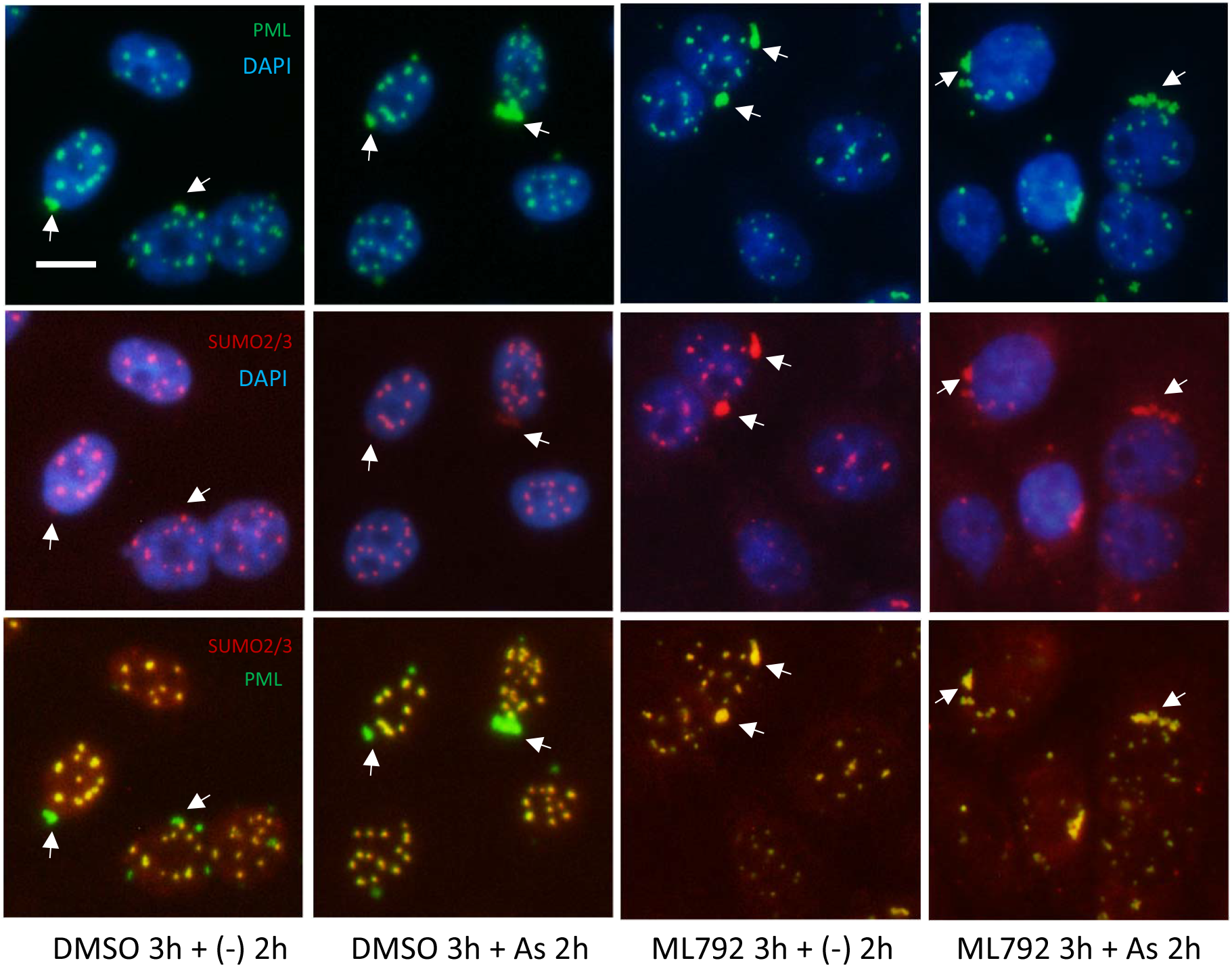
Fluorescent immunostaining of ML792-pretrreated CHOGFPPML cells to detect PML and SUMO2/3. The cells were treated with 0.1% DMSO or 20 μM ML792 for 3 h, then further treated with 3 μM As^3+^ or left untreated for 2 h. Note that SUMO2/3 in peri-nuclear PML aggregates (arrow) is less abundant in the ML792-treated cells compared to the DMSO-treated cells. Scale bar = 10 μm

## Discussion

### PML-NBs and peri-nuclear PML structures

Most PML-NBs occur in toroidal shapes in CHOGFPPML cells and in amorphous-like shapes in HEKGFPPML cells (Figs. 1 and 2). In addition to PML-NBs, peri-nuclear PML structures appeared to be accretions of several toroidal PML structures in CHOGFPPML cells. These observations suggest that the CHO cell has a propensity for annular PML-NBs, although it is not clear why the annular condensation of GFPPML occurs in CHO cells. Given that the SUMOylation of GFPPML proceeded normally in both cell types, PML-NBs may need some other scaffold or client molecules to form stable toroidal shapes. The disassembly of large toroidal PML-NBs to small dots in the HEKGFPPML cell nucleus (Fig. 2B) indicates that PML-NBs are not irreversible aggregates, and that toroidal PML-NBs in HEK cells are more labile than those in CHO cells. Irregular-shaped PML-NBs are prone to forming hydrogels with less fluidity in the sol-gel balance of non-membrane bodies because a liquid-like nature would make these bodies spherical due to viscous relaxation (32). Our current finding that toroidal PML-NBs were converted into small dots in the HEKGFPPML cell nucleus may shed light on the rheological nature of PML-NBs.

A systematic regulatory network deciphering study indicated that most cells with increased PML-NBs contain S phase DNA and low 5-ethynyl-2’-deoxyuridine (EdU) incorporation, suggesting that increased PML-NBs correlate with decreased DNA replication rates (33). It has been reported that the number of PML-NBs increases in early S phase in SK-N-SH and *GFP-PML-IV* transduced U-2OS cells due to fission of the parent NBs and not to increased PML protein levels (9). Although the transformation of large toroidal PML-NBs into small ones in HEKGFPPML cells (Fig. 2B) appeared to begin with dissolution or dissipation rather than fission in the present study, the intranuclear dynamics of PML-NBs may be fine-tuned with DNA synthesis.

### Effects of As^3+^, Sb^3+^, and Bi^3+^ on PML

The most notable biochemical changes in As3+-exposed cells was the solubility shift from the cold RIPA-soluble to the RIPA-insoluble form, and the SUMOylation of PML (34,35). These changes were also observed in Sb^3+^-exposed but not in Bi^3+^-exposed GFPPML-expressing cells (Fig. 3), consistent with findings using other types of cells (12,36). Bi^3+^ forms a stable complex with metallothionein in the cytosol and might not efficiently reach the nucleus (37). The solubility shift and SUMOylation of PML appear to be induced by TGF-β, as well as by As3+ or Sb^3+^ (38). The PML protein level, and the number and size of PML-NBs, are increased by IFNα in BL41 cells; however, the solubility change of PML does not occur in IFNα-treated cells. In contrast, TGF-β disrupts PML-NBs and changes PML from the RIPA-soluble (nucleoplasm) to the RIPA-insoluble form (nuclear matrix), and further enhances the SUMOylation of PML. Accordingly, a large amount of SUMOylated PML was found in the RIPA-insoluble fraction of TGF-β-treated cells when pre-treated with IFNα (38). The As3+-induced solubility shift is not limited to PML proteins. The majority of Hsp104p is recovered in the soluble fraction in untreated cells, whereas As3+ generates a concentration-dependent shift of Hsp104p from the soluble to the insoluble (aggregate) fraction in yeast (*Saccharomyces cerevisiae*) (39). The solubility shit and SUMOylation of PML in As3+-exposed cells are proposed to be mediated by binding of As3+ to cysteine residues in RBCC domain (11,13). The involvement of cytokines in As3+-exposed cells remains to be elucidated.

### A role of SUMOylation in PML-NBs

MAPPs are aggregated forms of PML that appear after nuclear membrane breakdown, and CyPNs are formed in the peri-nuclear regions of daughter cells after mitosis. More MAPPs or CyPNs remain in the cells if the recycling of PML is retarded by As^3+^-treatment (40). MAPPs and CyPNs appear to be devoid of SUMO, Sp100, and Daxx, which are well-known PML-NB client proteins (41,42). We did not differentiate MAPPs and CyPNs, referring to these PML aggregates as extranuclear or peri-nuclear PML structures in the present study. Although the biological functions of these peri-nuclear PML structures have not been elucidated (43), these aggregates can be used as markers of laminopathies (44). Our findings that PML colocalizes with SUMO2/3 in PML-NBs, and that peri-nuclear PML structures contain less SUMO2/3, are consistent with these previous studies. SUMO2/3 distributed almost evenly between PML-NBs and peri-nuclear PML structures in the presence of ML792, an inhibitor of SUMO-activating enzymes, irrespective of As^3+^ exposure (Fig. 10). Since the SUMOylation of GFPPML was almost completely inhibited by ML792 (Fig. 9), it is probable that SUMO monomers non-covalently associate with peri-nuclear PML structures as well as with PML-NBs in the presence of ML792.

The 69-kDa isoform of PML mutated at the B-box (B1C17C20AΔAla, B1C25C28AΔAla, and B2C21C24ΔAla) does not form PML-NBs and is instead present diffusely over the nucleus (10). The three main SUMOylation sites (K65, 160, and 490) and the SIM are not essential for formation of the PML multimer lattice or the outer shell of PML-NBs. SUMO2/3 appears to be more closely associated with PML compared to other partner proteins such as Sp100, Daxx, SUMO1, and RNF4 (45). In addition, depletion of SUMO3 reduces the number of PML-NBs (46). Recently, PML-specific sequences (F52QF54 and L73) were shown to play essential roles in PML-NB assembly, recruitment of Ubc9, and SUMOylation (14). The depletion of SUMO-specific protease 1 (SUSP1/SENP6), which dismantles highly conjugated SUMO2/3 proteins, caused marked accumulation of SUMO2/3 in PML-NBs (47) and increased the average number of PML-NBs per nucleus from 3.7 to 5.9 in HeLa cells (48). Thus, the balance of SUMOylation and deSUMOylation of PML is critical for the appropriate maintenance of PML-NBs. In addition to the SUMOylation and deSUMOylation of PML, PML-NBs might be regulated by non-PML-NB-associated proteins. The expression of either Ki-1/57 or CGI-55 proteins, which have three SUMOylation sites, reduces the number of PML-NBs in both untreated and As^3+^-treated HeLa cells, whereas the expression of Ki-1/57 mutant (K/3R) reduces the number of PML-NBs in untreated cells but not in As^3+^-treated cells (49). The inhibition of SUMOylation by ML792 increased the size of PML-NBs in the present study (Fig. 8), suggesting that SUMOylation indeed regulates the scaffold structure of PML-NBs.

Blocking ubiquitination using TAK243 resulted in the build-up of SUMOylated proteins in HeLa and U-2OS cells, with PML-NBs apparently functioning as primary accumulation sites for newly synthesized and SUMOylated proteins (30). Ubiquitination by RNF4 E3 ligase via reciprocal SUMO-SIM interaction is proposed to be the main cascade for the proteasomal degradation of SUMOylated PML-RARα (7,17,50). PML-NBs may be transiently employed to store harmful SUMOylated proteins by phase separation before proteolysis by the ubiquitin-proteasome system (51). However, functional roles of SUMOylated PML are not clear, because modification with SUMO protects IκBα (52) and Glis2/NPHP7, a Cys2/His2 zinc finger transcription factor (53), from ubiquitin-proteasome degradation by competing the same lysine (2). An ancillary but interesting finding in the present study is that the cytotoxic effect of TAK243 was antagonized by ML792 (Supporting Fig. 2), suggesting that SUMOylation may be harmful to the cells when ubiquitination is inhibited. In summary, the shapes of PML-NBs and extranuclear PML structures depend on the cell type. As^3+^ and Sb^3+^ cause similar biochemical effects on PML and agglomerate of PML-NBs. SUMOylation of PML regulates the size and number of PML-NBs.

## Experimental procedures

### Chemicals

TAK243 (Ub-activating E1 enzyme inhibitor, Selleckchem, Houston, TX) and ML792 (SUMO E1 inhibitor, MedKoo Bioscience, Morrisville, NC) were dissolved in DMSO and used at a final concentration of 0.1% DMSO. A 20S proteasome assay kit was purchased from (Enzo, Famingdale, NY). Bismuth (III) nitrate pentahydrate and antimony (III) chloride were purchased from Wako (Osaka, Japan). They were dissolved in 0.1 M citrate buffer (pH 7.4) at 10 mM and used as stock solutions. Sodium *m*-arsenite and purchased from Sigma-Aldrich (St. Louis, MO) and dissolved in PBS solution. These solutions were sterilized using a syringe filter (0.45 μm pore size). Unless otherwise specified, common chemicals of analytical grade were obtained from Sigma-Aldrich or Wako.

### Cells and GFP monitoring with confocal microscopy

The human *PMLVI* gene engineered into the pcDNA6.2N-EmGFP DEST vector was transduced into HEK293 and CHO-K1 cells (54). Stable transfectants were selected using blasticidine S and designated as HEKGFPPML and CHOGFPPML cells, respectively. The cells were cultured in glass-bottom culture dishes and GFP fluorescence was monitored by confocal lase microscopy (TCS-SP5, Leica Microsystems, Solms, Germany) or fluorescence microscopy (Eclipse TS100, Nikon, Kanagawa, Japan).

### Protein extraction and western blot

The cell monolayers were rinsed with Hank’s balanced salt solution (HBSS) and lysed with ice-cold RIPA buffer (Santa Cruz, Dallas, TX) containing protease (Santa Cruz) and phosphatase inhibitor cocktails (Calbiochem/Merk Millipore, San Diego, CA) on ice for 10 min. The lysate was centrifuged at 9,000 *g* for 5 min at 4 °C. The supernatant was collected and labelled as the RIPA-soluble fraction (Sol). The pellet, which contained nucleic acids, was rinsed with PBS and treated with Benzonase^®^ nuclease (250 U/mL in Tris buffer, Santa Cruz) of the same volume as the RIPA buffer at 25 °C for 2 h with intermittent mixing (Thermomixer comfort, Eppendorf, Wesseling-Berzdorf, Germany). The digested sample was labelled as the RIPA-insoluble fraction (Ins). The following antibodies were used for immunoblot analyses, with dilution rates of 1:750 for primary antibodies and 1:2500 for secondary antibodies in Can-Get-Signal^TM^ solution (TOYOBO, Osaka, Japan): anti-PML antibody (Bethyl, A301-167A, rabbit polyclonal); anti-SUMO2/3 (MBL, 1E7, mouse, monoclonal); anti-multi ubiquitin (MBL, FK2, mouse, monoclonal); anti-SUMO1 (CST, C9H1, rabbit, monoclonal); anti-UBA2 (CST, D15C11, rabbit, monoclonal); anti-GFP (Abcam, goat, polyclonal); HRP-conjugated anti-histone H3 (CST, #12648, rabbit, monoclonal); HRP-conjugated anti-α-tubulin (MBL, #PM054-7, rabbit, polyclonal); HRP-conjugated goat anti-rabbit IgG (Santa Cruz, sc-2054); HRP-conjugated goat anti-mouse IgG (Santa Cruz, sc-2055); HRP-conjugated mouse anti-goat IgG (Santa Cruz, sc-2354). Aliquots of the RIPA-soluble fraction and the nuclease-treated pellet suspension (RIPA-insoluble fraction) were mixed with LDS sample buffer (1x TBS, 10% glycerol, 0.015% EDTA, 50 mM DTT, and 2% LDS) and heated 95 °C for 5 min. Proteins in the samples were resolved by LDS (SDS)-PAGE (4-12%) and electroblotted onto PVDF membranes. The membranes were blocked with PVDF Blocking Reagent (TOYOBO) before probing with antibodies. Signals on the ECL (Prime, GE Healthcare, Buckinghamshire, UK)-treated PVDF membrane were then captured with a CCD camera (Lumino Imaging Analyzer, FAS-1100, TOYOBO; Amersham ImageQuant 800, GE Healthcare, Uppsala, Sweden).

### Immunofluorescent staining

The cells were grown in an 8-well chamber slide (Millicell EX slide, Merck Millipore, Burlington, MA) to early confluency. For HEKGFPPML cells, the chamber slide was treated overnight with 50 μg/mL collagen Type IV solution (Wako). The cells were washed with warmed (37 °C) HBSS and fixed with 3.7% formaldehyde solution for 10 min, permeabilized with 0.1% Triton X-100 for 10 min, and treated with Image-iT™ FX Signal Enhancer (Invitrogen-ThremoFischer, Carlsbad, CA) for another 10 min. The cells were immunostained with Alexa Fluor^®^ 594-labeled anti-SUMO-2/3 for 45 min. Anti-SUMO2/3 antibody (MBL) was labelled with Alexa Fluor^®^ 594 using an Alexa Fluor™ 594 Antibody Labeling Kit (Invitrogen-ThermoFischer) and diluted to 1/200 in Can-Get-Signal^TM^ Immunostain solutions (TOYOBO). DAPI was used to counter-stain the nuclei. The fluorescent images were captured using a fluorescence microscope (Eclipse 80i, Nikon, Tokyo) and the digital images were assembled using Adobe PHOTOSHOP^®^ software.

### Immunoprecipitation

The cells were lysed with cold RIPA solution and clear supernatants were obtained by centrifugation (9000 *g*, 5 min). The protein concentration of the supernatant was adjusted to 2 mg/mL with RIPA. The sample was diluted to one fourth the initial concentration with 150 mM Tris-HCl solution (pH 7.2) and reacted with anti-GFP mAb-magnetic beads (MBL) or control IgG2a-magenitic beads (MBL) at 4-8 °C for 2 h with intermittent vortexing. The magnetic beads were washed 3 times with 150 mM Tris-HCl solution (pH 7.2) containing 0.1% NP40 and the pellet was boiled with 2X LDS(SDS)-PAGE sample buffer for 5 min. The sample was centrifuged, then the supernatant was subjected to LDS (SDS)-PAGE and subsequent western blot analysis as described above.

### Cell viability

The cells were pre-cultured to early confluency in a 96-well culture dish, then further cultured in the presence or absence of As^3+^, Sb^3+^, Bi^3+^, or ML792 and/or TAK792. The cells were washed twice with HBSS and the number of viable cells was assayed colorimetrically by WST-8 (Wako) using a microplate reader (POLARstar OPTIMA, BMG Labtech, Offenburg, Germany).

### Data analyses

Data are presented as means ± SEM. Statistical analyses were performed by ANOVA followed by Tukey’s post-hoc test. A probability value of less than 0.05 was accepted as indicative of statistical significance. The number and size analyses of PML-NBs were performed using ImageJ (NIH, https://imagej.nih.gov/ij/).

## Acknowledgment

The authors would like to thank Dr. Ayaka Kato-Udagawa and Ms. Mihoko Tadano for their technical assistance.

## Author contributions

SH designed the study, performed the experiments, and wrote the paper. OU performed the experiments and wrote the paper.

## Funding

This work was partially supported by a Grant-in-Aid from the Japan Society for the Promotion of Science (16K15386).

## Conflict of Interest

The author has no conflicts of interest regarding the contents of this article.

---

**Supporting Fig. 1A.**
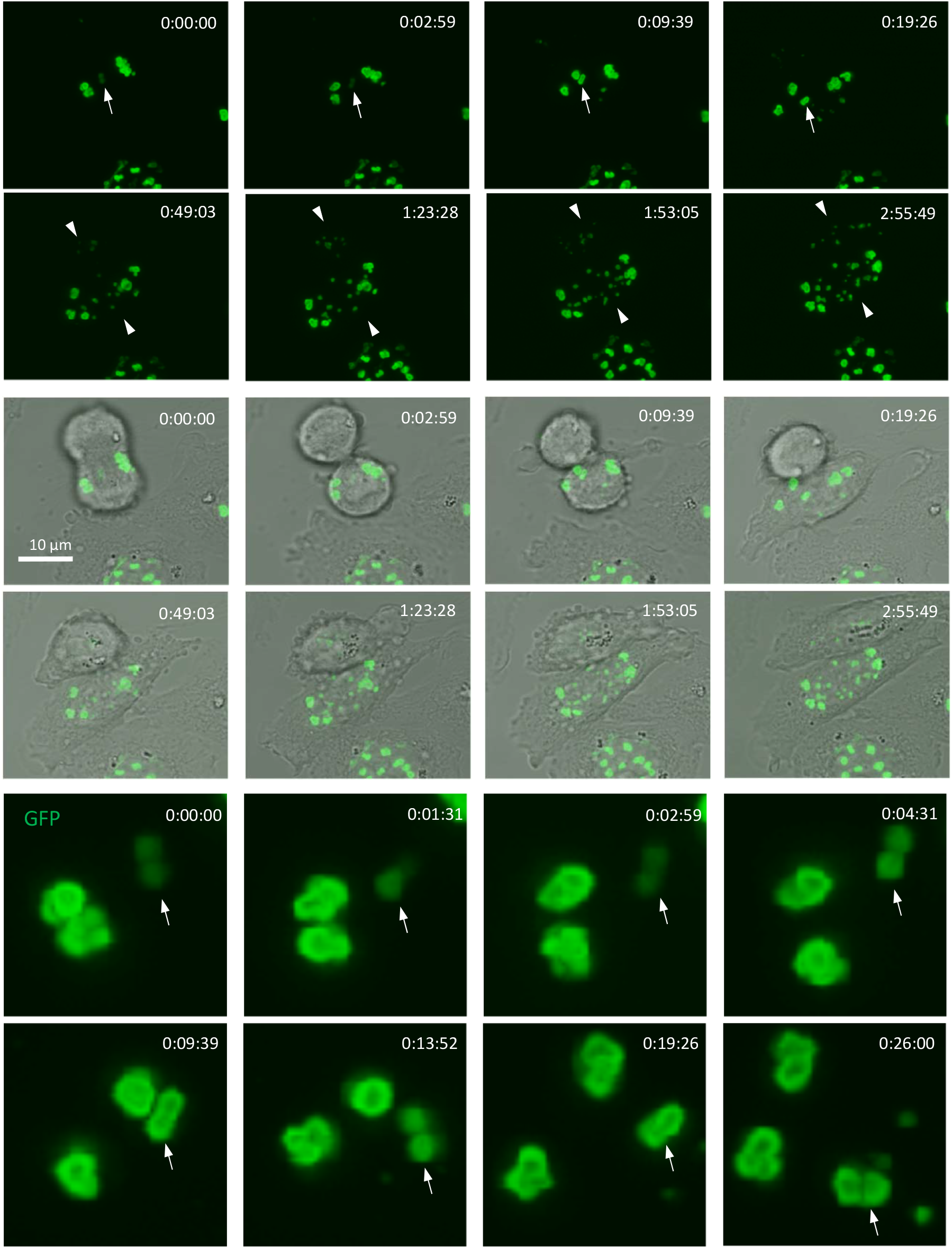
Emergence and uneven partitioning of peri-nuclear PML structures in dividing untreated CHOGFPPML cells. Live GFP images were captured during cell division by confocal laser microscopy in z-stacking mode. The upper and middle 8 panels show GFP alone and the corresponding GFP-bright field overlaid images, respectively. The lower 8 panels show enlarged and intensified GFP images of a peri-nuclear PML structure indicated with arrows on the top panels with four additional time-lapsed GFP images. The time counter is shown in the right upper corner of each panel. The peri-nuclear PML structures appear to be comprised of 2–4 toroids. The nascent small PML-NBs (closed arrowheads) appeared as the daughter cells spread.

**Supporting Fig. 1B.**
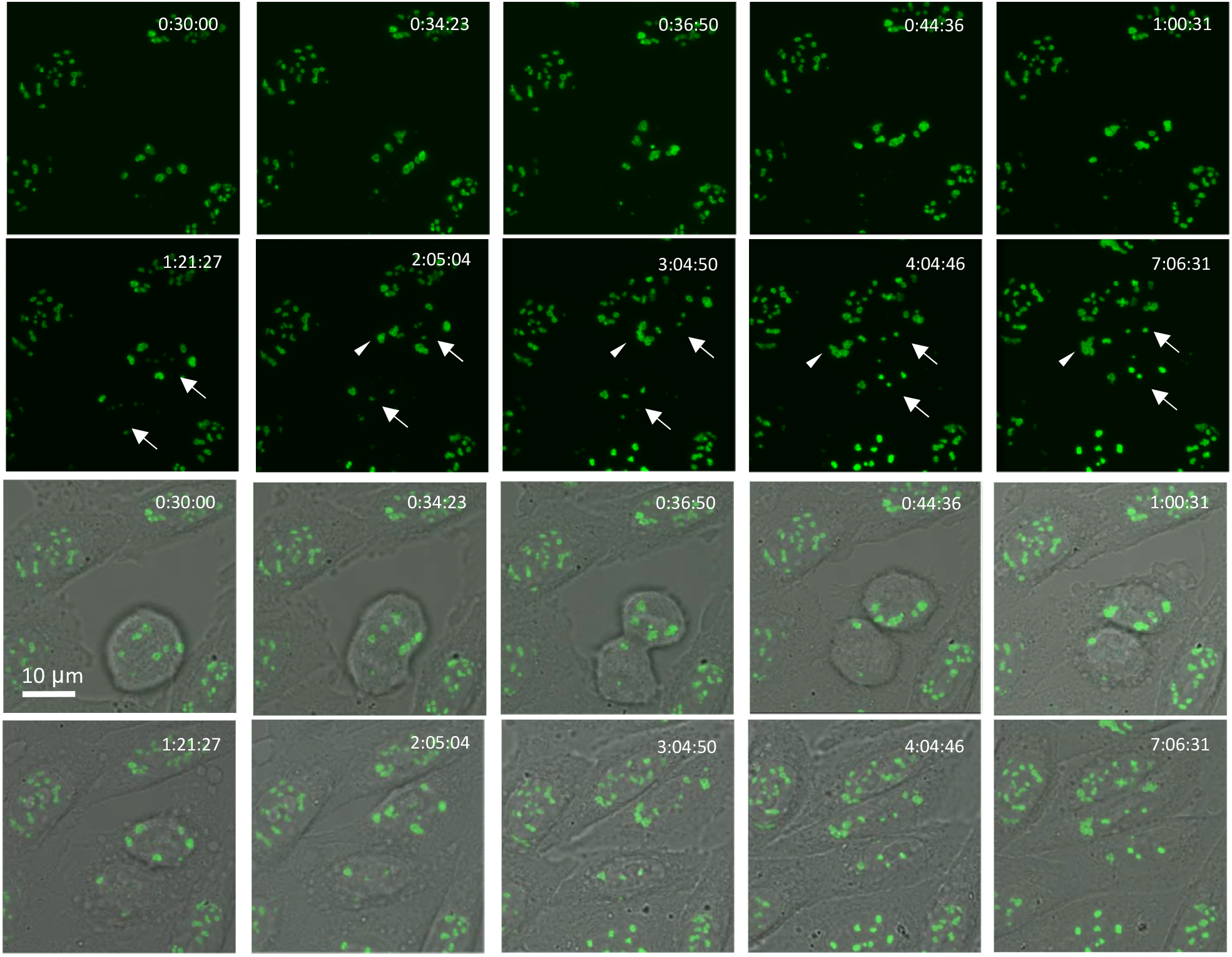
Uneven partitioning of peri-nuclear PML structures in As^3+^-exposed CHOGFPPML cells. Image capture started 30 min after the addition of As^3+^. The upper and lower 10 panels show GFP images alone and the corresponding GFP-bright field overlaid images, respectively. See also the legend to Supporting Fig. 1A. The nascent small PML-NBs (arrowheads) appeared as the daughter cells spread, regardless of As^3+^ exposure.

**Supporting Fig. 2.**
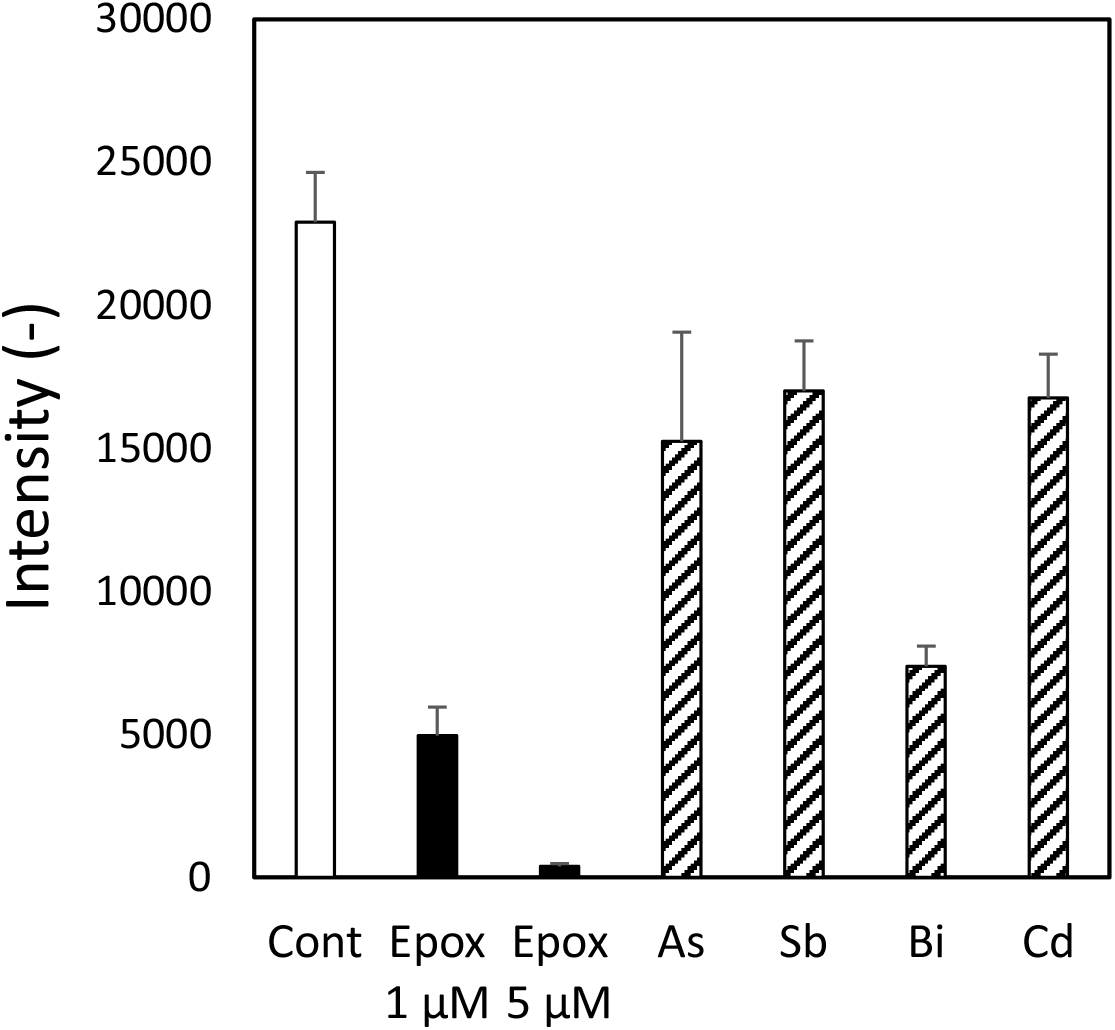
Inhibition of proteasomal activity by As^3+^, Sb^3+^, Bi^3+^, and Cd^2+^. The proteasomal inhibitory effect was assayed using a 20S proteasome assay kit. Briefly, the fluorescence (Ex/Em, 360/460 nm) of the reaction mixture (proteasome, Suc-LLVY-AMC, and SDS) was monitored every 1 min in the presence or absence of As^3+^, Sb^3+^, Bi^3+^, Cd^2+^, or epoxomicin (a positive control) at 30oC for 20 min using a microplate reader (Infinite M Plex, TECAN, Männedorf, Switzerland). As^3+^, Sb^3+^, Bi^3+^, and Cd^2+^ were added to the reaction mixture at a final concentration of 30 μM. The fluorescence was increased linearly and the increment of fluorescence in 20 min was used for evaluation of relative inhibitory activity of proteasome. Data was presented as mean ± SEM of 4 replicate measurements.

**Supporting Fig. 3.**
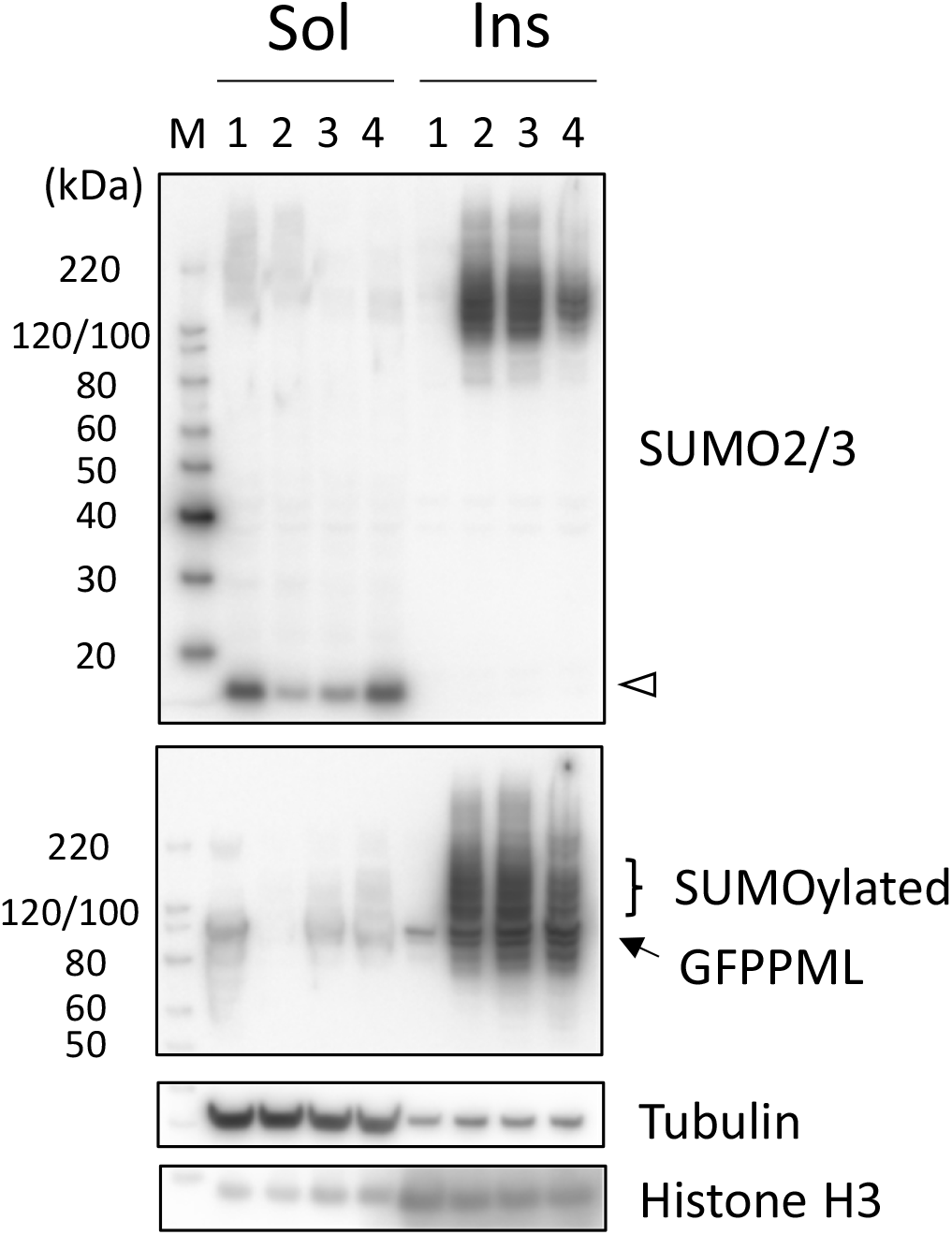
Recovery of SUMO2/3 and GFPPML proteins in the RIPA-soluble fraction (Sol) from 24 h-exposure to As^3+^ in HEKGFPPML cells. The cells were exposed to 3 μM As^3+^ for 24 h or left untreated. The As^3+^-exposed cells were lysed with the RIPA buffer immediately or washed and further cultured in As^3+^-free culture medium further for 8 or 24 h. 1, untreated; 2, 24 h exposure to As^3+^; 3, 24 h exposure to As^3+^and 8 h recovery in As^3+^-free culture medium; 4, 24 h exposure to As^3+^and 24 h recovery in As^3+^-free culture medium. An open arrowhead indicates SUMO2/3 monomers. The unconjugated GFPPML and GFPPML conjugated with SUMO2/3 in the RIPA-insoluble fraction (Ins) decreased during the culture in As^3+^-free culture medium.

**Supporting Fig. 4.**
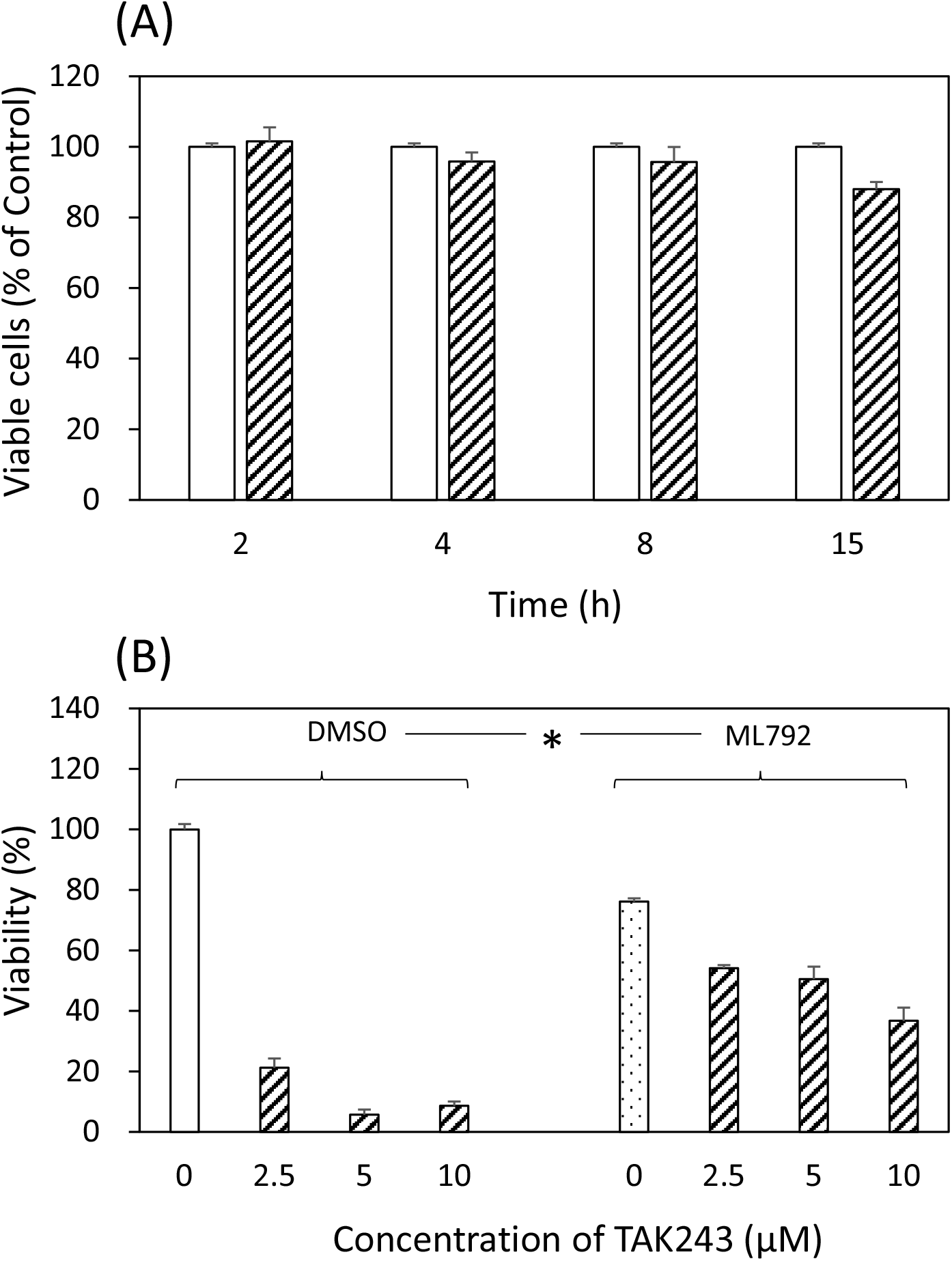
Time-course changes in the number of viable ML792-exposed cells (A) and the inhibitory effects of ML792 on the cytotoxicity of TAK243 (B) in HEKGFPPML cells. (A) The cells were exposed to 0.1% DMSO (open columns) or 20 μM ML792 (hatched columns) for 2, 4, 8, 15 h. The number of viable cells was assayed using WST-8 agent. Data are presented as means ± SEM of five wells. (B) The cells were exposed to 0, 2.5, 5, and 10 μM TAK243 for 24 h in the presence or absence of 20 μM ML792. Data were analyzed by two-way ANOVA followed by Tukey’s multiple comparison. The viability of ML792-treated cells was significantly elevated compared to 0.1% DMSO-treated cells in the presence of 10 μM TAK243.

